# Meta2Data: Automated data processing for large scale microbiome data re-analysis

**DOI:** 10.64898/2026.10.01.732036

**Authors:** Jiaxuan Li, Anne Duplouy, Linyang Sun

## Abstract

Meta2Data is a modular, largely automated pipeline implemented in Python and Bash for the retrieval, standardization, and processing of public microbiome sequencing data. It retrieves metadata from major public repositories, including INSDC and CNCB, using user-provided keywords or BioProject/BioSample identifiers. Retrieved metadata are standardized using a synonym dictionary and then subjected to manual curation. Based on the curated metadata, Meta2Data downloads the corresponding raw sequencing data and processes them using standardized workflows based on DADA2 or VSEARCH. The pipeline supports both single- and multiple-target-region datasets and produces integrated feature tables, taxonomic assignments, and phylogenetic trees for downstream microbial community analyses.

**Availability and Implementation:** Meta2Data is freely available at https://github.com/LinyangSun/Meta2Data and archived on Zenodo at https://doi.org/10.5281/zenodo.20670433

## Introduction

The volume of data available from public data repositories is rapidly growing (Gaffney, et al., 2020). The accumulation of metabarcoding data from worldwide projects and organisms, which makes broad-scale re-analyses of global microbiome compositional patterns (Cheng, et al., 2025; Li, et al., 2023; Thompson, et al., 2017), species interaction networks (Lam and Ye, 2022), or functionality (Adejumo, et al., 2026; Hoffbeck, et al., 2025; Wirbel, et al., 2019), increasingly feasible (Gao, et al., 2023; Katz, et al., 2022).

The re-analyses, however, remain labor-intensive due to the large volume of data they include. Additionally, the acquisition and re-use of such deposited data are challenged by inconsistent metadata annotations between repositories, or inconsistent or incomplete application of standardized repository practices (e.g., missing primer sequences). To date, existing data retrieval tools mostly, target databases within the International Nucleotide Sequence Database Collaboration (INSDC), often overlooking other public repositories such as the China National Center for Bioinformation (CNCB). Moreover, most available analytic tools cannot automatically analyze datasets sequenced on different platforms or derived from different hypervariable regions. Such constraints reduce the size and breadth of the datasets available, limiting the analytical power and comprehensiveness of microbiome re-analysing studies.

Despite the recent development of tools that can address some of these challenges individually, including metadata retrieval (Eaton, 2020; Fumagalli, et al., 2023; Gálvez-Merchán, et al., 2023; Woodcroft, et al., 2024) and curation (Abdill, et al., 2025; Kasmanas, et al., 2021; Zhang, et al., 2026; Ziemski, et al., 2022), no single tool yet integrates all steps within a unified workflow. Here, we present Meta2Data, a largely automated, modular pipeline that can complete metadata retrieval and cleaning, as well as sequence retrieval and processing. The pipeline can process short-read and long-read amplicon sequencing data using different denoising methods (DADA2 or Vsearch), and can produces phylogenetic and taxonomy outputs.

## Implementation

Meta2Data operates through a series of automated steps, organized into three parts: metadata retrieval and pre-cleaning (MetaDL), sequence data preprocessing (AmpliconPIP), and taxonomic annotation and phylogenetic tree construction (AmpliconTAXA) (Figure 1). Meta2Data was written by Python and Bash, can be run on Linux, and provides a SIF container version. The source code and SIF container are available at: https://github.com/LinyangSun/Meta2Data/pkgs/container/meta2data.

**Figure 1.**
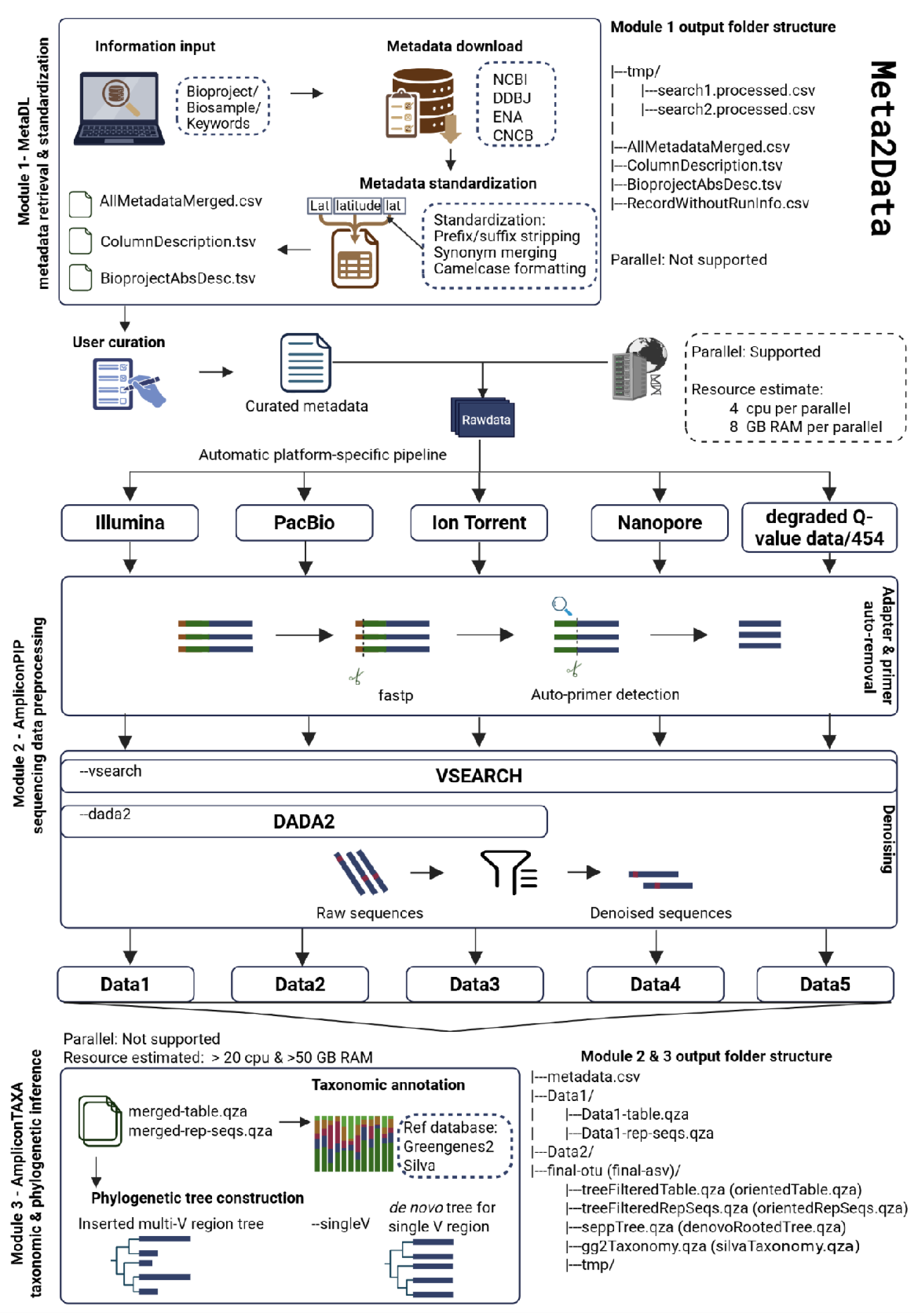
Overview of the Meta2Data pipeline. First, the MetaDL module retrieves and homogenizes the metadata from public repositories (NCBI, DDBJ, ENA, CNCB). Then, the AmpliconPIP module routes the raw sequence data to platform-specific denoising pathways. Under ‘--dada2’ mode, data produced on Illumina, PacBio, and Ion Torrent sequencing platforms are processed with DADA2, while quality-degraded datasets are rejected; under ‘--vsearch’ mode, datasets from all platforms are uniformly processed with VSEARCH. Adapter and primer sequences are automatically detected and removed prior to denoising. Within the AmpliconTAXA module, the sequences are merged, and taxonomic annotations against reference databases are performed before constructing a final phylogenetic insertion tree. Output folder structures and resource estimates for each module are shown alongside the workflow.

### 1) MetaDL: Metadata and sequence acquisition

The module MetaDL queries both INSDC and CNCB repositories by directly using user-provided keywords, BioProject or BioSample IDs. The retrieved metadata may contain inconsistent or non-standardized attribute names. MetaDL standardizes these attributes using a dictionary of synonymous attribute names, which is a combination of official dictionaries (https://www.ncbi.nlm.nih.gov/biosample/docs/attributes/ and https://ngdc.cncb.ac.cn/gsub/document/Handbook-BioSample.pdf) and groups of similar attributes derived from publicly available NCBI BioSample and SRA metadata. For example, the synonymous attributes ‘latlon’, ‘lat lon’, or ‘lat_lon’ are all clustered under the standardized key ‘LatitudeAndLongitude’. When multiple synonyms map to the same standardized attribute, MetaDL compares their values row by row, and only merges consistent synonyms. If any conflicting non-missing values are found, original attribute names and values are kept. MetaDL standardizes attributes only, and does not perform automatic splitting or hierarchical inference. Detailed methods are provided in Supplementary file (Figure S1).

MetaDL outputs a harmonized sample-level metadata table, a summary of each attribute’s fill-rate and value, and a BioProject-level description file containing published projects information. These three outputs can be manually curated by the user, who can further review and select the final dataset to be used in the following sequence-processing steps. The user then passes the curated metadata to the AmpliconPIP module for automated sequence acquisition by simply providing the Bioproject and SRA ID column names.

### 2) AmpliconPIP: Automatic primer detection and Data pre-processing

The presence of erroneous primer information in metadata is one issue of public microbiota data processing. The AmpliconPIP module addresses this issue using an automated primer detection approach (APD), which detects and removes primers even when primer information in the metadata is missing or incorrect. The APD approach examines the beginning of reads, identifies sequence patterns characteristic of primers, and compares these patterns to known primer sequences. When no known primer is identified, APD uses sequence degeneracy to distinguish a potential unknown primer from natural biological variation (Fulghum, et al., 2024; Li, et al., 2012). Each step of the APD approach is described briefly below, and further details on a benchmark design, parameter evaluation, and results of a mock-community analysis are provided in Supplementary files (Figure S2-3).

First, the APD approach removes potential adapters using fastp (Chen, et al., 2018). Second, nucleotide frequencies are calculated for each of the first 20 base positions across all sequence-reads from each project. Nucleotide base frequencies ≥10% are retained, and detected bases at each position are assigned to either Conserved C=1, Double D=2, Triple T=3, or Quadruple Q=4. These values are used to construct a consensus sequence and to calculate its fold degeneracy across the 20 positions. The consensus sequence is then compared against a built-in primer motif database generated from consensus primer regions. If a match is found, the primer sequence is trimmed. Otherwise, the calculated fold degeneracy is used to distinguish any potential unknown primer from a natural sequence variation (Rose, et al., 2003; Rose, et al., 1998). Sequences with a fold degeneracy <16 are classified as unknown primers and the primer regions are then trimmed, whereas sequences with a fold degeneracy ≥16 are treated as natural sequence variation, and conserved. Alternatively, users can specify the ‘--skip-unknown-primers’ parameter to skip datasets from further processing unknown primer regions. To demonstrate the performance of this APD approach and its parameter thresholds, we performed two benchmark analyses, which showed reliable APD performances, with only error rates increasing in extreme low diversity datasets, containing 1–5 unique sequences.

After primer detection and trimming, the sequence datasets are routed to either DADA2- or VSEARCH-based sequence-denoising and -processing workflow. The ‘--dada2’ mode uses error-model-based sequence inference and therefore only process sequences with reliable per-base quality scores. In contrast, the ‘--vsearch’ mode uses clustering-based sequence processing and does not rely on an explicit sequencing-error model, making it suitable for datasets of sequences with degraded or unreliable Phred quality scores. Under the ‘--dada2’ mode, Illumina datasets with reliable quality scores, PacBio and Ion Torrent datasets are accepted; otherwise, datasets are skipped. Under the ‘--vsearch’ mode, datasets from all sequencing platforms should be accepted. All datasets rejected during preprocessing are reported in the output.

### 3) AmpliconTAXA: Cross-dataset taxonomy harmonization and phylogenetic integration

Integrating data from different sequencing regions, particularly for phylogenetic reconstruction, has remained challenging. To address this, the AmpliconTAXA module orientates all sequences against a uniform strand from the Greengenes2 2024.09 sequence orientation reference (McDonald, et al., 2024) using RESCRIPt (Robeson, et al., 2021). Sequences that cannot be oriented or aligned to the reference are filtered out, as without this orientation step, the accuracy of the next step will drop.

Taxonomy is then assigned to the sequences using a user-selected pre-trained Naive Bayes classifier, such as Greengenes2 2024.09 (McDonald, et al., 2024)or SILVA 138.2 (Chuvochina, et al., 2026). Meta2Data provides two options for phylogenetic tree construction. In the default mode, which accommodates sequences from multiple sequencing regions, sequences are integrated into a full-length 16S reference tree using the fragment-insertion (SEPP) (Janssen, et al., 2018) function in QIIME 2. Sequences that cannot be placed on the reference tree are discarded from the output count table and the representative-sequences file. When the option ‘--singleV’ is enabled, the phylogenetic tree is constructed *de novo* from the representative sequences. Phylogenetic tree construction can be omitted by specifying the ‘--notree’ option. We performed a Benchmark 3 to evaluate the robustness of this workflow (Supplementary File; Figure S4).

### 4) Calculation resource, reproducibility, and transparency

The pipeline generates log files and summary statistics throughout the analyses, and the ‘--parameter’ option allows users to modify selected parameters as needed. The AmpliconPIP module supports parallel processing of diverse data. After assessing the competence of this module in a Benchmark 4 (Supplementary File; Figure S5), we recommend that the user secures at least 4 CPUs and 8 GB memory per process, with total resources scaled by the number of concurrent processes. For the AmpliconTAXA module, the SEPP-based tree construction requires substantial resources, and we recommend that users secure 20 CPUs and at least 60 GB of memory, with longer wall time for larger datasets, before launching this module. Omitting tree construction would reduce runtime and CPU use.

## Discussion

Meta2Data is designed to optimize the efficiency and quality of data retrieval for meta-analyses of metabarcoding data (Raassina, et al., 2026). We describe below practical considerations that should be noted before using the pipeline:

- The Meta2Data pipeline still includes one necessary manual step, during which the user must review the extracted dataset. At this stage, the user evaluates the quality of the dataset using the summary file generated, and identify whether some data requires to be reprocessed or excluded.
- Cross-study integration can introduce batch effects arising from differences in sequencing platforms, targeted hypervariable regions, and primers. Meta2Data can reduce variation introduced by inconsistent processing workflows, but at no point the pipeline corrects for study-specific batch effects. Users must evaluate and account for these effects in their own downstream analyses.
- Finally, Meta2Data produces standard QIIME 2 outputs, including feature tables, taxonomy tables, and rooted phylogenies. Although these outputs can be used in QIIME 2 or R for downstream analyses, such as diversity analysis and statistical modelling, these analyses are currently beyond the scope of the Meta2Data pipeline.

## Acknowledgement

We thank the Research Council of Finland (grants # 321543 and #355152 to AD) for financial support and IT Center for Science (CSC), Finland, for providing computational resources. We thank Dr. Claire Williams and Tina Raassina, early testers of the pipeline, for providing useful feedback.

