## Supplementary file for "Meta2Data: Automated data processing for large scale microbiome data re-analysis"

Supplementary Methods

1. Expanding the attribute dictionary

Metadata records from public samples stored in NCBI (https://ftp.ncbi.nlm.nih.gov/) were used as the source data of a broad dictionary of attribute labels/terms.

We used BioSample and SRA metadata in the following tests. For each attribute, we sampled 32 random combinations of values and their associated units, excluding any sample with missing values. These combinations were used to compare the information recorded under different attributes using a schema matching approach . The comparisons were performed using the COMA algorithm implemented in the software framework/tool Valentine (Koutras, et al., 2021). Results from these comparisons identified attributes that may describe similar information, which we subsequently clustered into distinct attribute groups. We then ranked the attribute labels (i.e., terms) from each attribute group based on how often they occurred in each source, and selected those accounting for 90% of all label occurrences. In total, 245 and 337 attribute groups accounted for 90% of the label occurrences from BioSample and SRA, respectively. Among them, 238 attribute groups overlapped between the two metadata sources, leaving a total of 344 unique attribute groups after combination of the two sources (Figure S1). Every member of a group containing at least one of these high-frequency attributes was retained.

All attribute groups were then merged into our built-in dictionary. If a group matched an already existing dictionary entry, any new terms were added to their corresponding entry in the dictionary. Groups with no existing match in the dictionary were simply added as new key-value pairs to the dictionary. We normalized the spelling/format of each attribute label using camelCase normalization. If the camelCase forms of all attributes were identical, their corresponding groups were merged into the dictionary using the camelCase format of the key. Groups containing inconsistent camelCase formats were discarded.

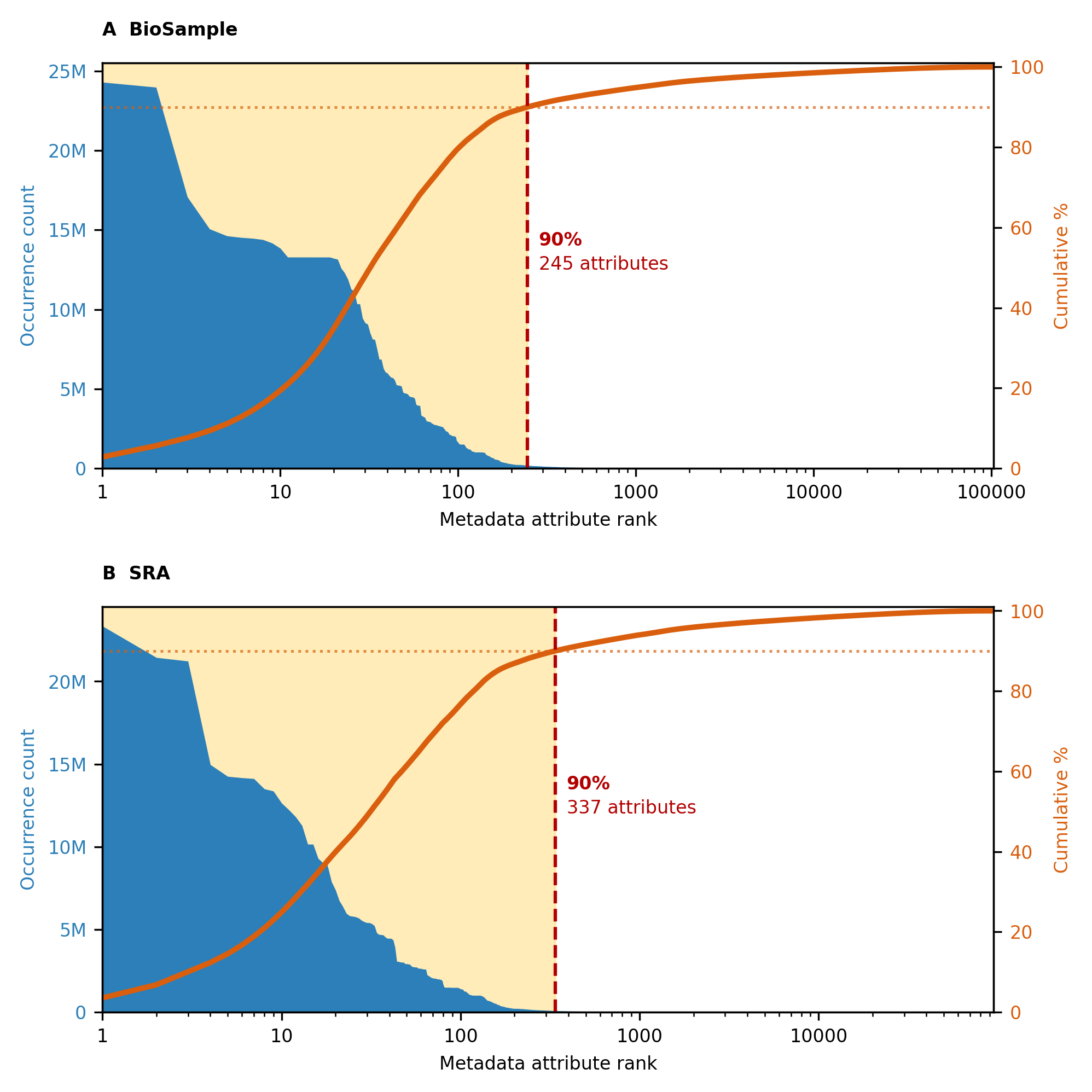

**Figure S1. Frequency distribution of metadata attributes in BioSample (A) and SRA (B).** Attributes are ranked by decreasing occurrence on a log scale. Blue areas show occurrence counts (left axis), and orange curves indicate cumulative percentages of frequency (right axis). Yellow shading and red dashed lines mark the top 245 BioSample attributes and the top 337 SRA attributes that together account for 90% of attribute occurrences in each source.

2. Benchmark 1: Testing for incorrect primer trimming

2.1 Methods

A collection of 390,271 distinct bacterial and archaeal 16S reference sequences from SILVA 138.2, and 22 primers, were used to test the efficiency of Meta2Data’s primer trimming approach. To identify the binding site of each primer, we took the 30 bases after each primer. To identify the best parameters for the primer detection workflow, we also test the same libraries using primer length windows covering the first 15, 20, or 25 bases of each sequence, and six different fold thresholds (i.e., 4, 8, 16, 32, 64, and 128). We then built a total of 30 libraries for each primer tested (30x22=660), every three replicated libraries contained 10,000 reads including either 1, 2, 3, 5, 10, 20, 50, 100, 200, or 500 unique sequences. Unfortunately, six libraries could not accumulate sufficient unique sequence (richness=500), and were excluded, leaving 654 libraries for the analysis.

Benchmark 1 evaluated false trimming by Meta2Data using 654 simulated libraries, from which primers had already been removed. Three detection windows (15, 20, and 25 bp) and six different fold-thresholds (4, 8, 16, 32, 64, and 128) were tested, yielding 18 parameter combinations, and a total of 11,772 evaluations.

2.2 Results

Because the expected trimming length was zero for all libraries, the false-trimming rate was defined as the proportion of libraries with a predicted trimming length greater than zero. Our analysis revealed that false trimming occurs predominantly in libraries with low sequence richness. At richness levels of 1–5, the false-trimming rate varied from 100% to 4.55%. No false trimming was observed in libraries including >5 unique sequences, when the fold threshold was below 16. These results suggest poor performance of our trimming workflow only in samples with extremely low sequence-diversity. Increasing the window length reduced the error rate, whereas increasing the fold threshold increased the error rate (Figure S2).

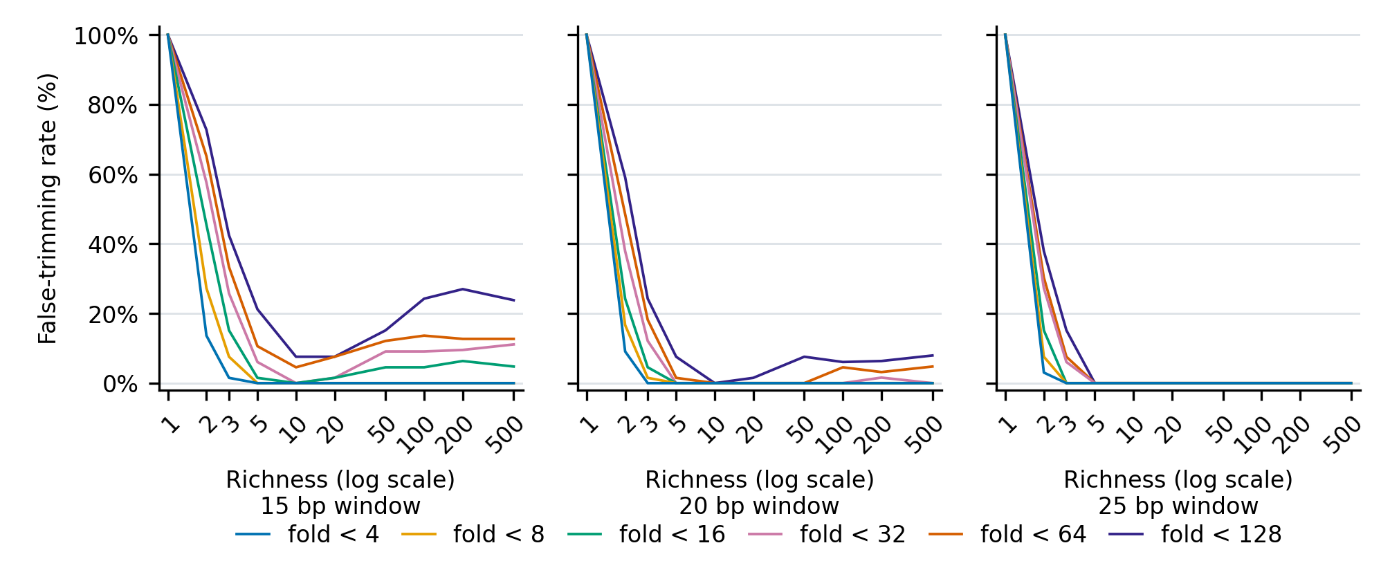
**Figure S2:** **False primer-trimming rates across libraries with varying levels of unique sequence richness.** Panels show results for target sequence lengths of 15 bp (left), 20 bp (middle), and 25 bp (right). Colored lines indicate different fold thresholds (4, 8, 16, 32, 64, and 128). Richness is plotted on a logarithmic scale, and false primer-trimming rates are presented as percentages (%). False primer-trimming rate was calculated as the percentage of reads removed from primer-trimmed libraries.

3. Benchmark 2: Testing the detection and successful trimming of external primers

3.1 Methods

To construct Benchmark 2, we used the same SILVA reference sequences as in Benchmark 1. We extracted 30 nucleotides from the start of each primer-binding site and set the target richness to 500 distinct sequences per library. Primer sequences were collected from PIMENTO, ProbeBase, and published studies (Chen et al., 2018; Greuter et al., 2016; Klindworth et al., 2013; Regueira-Iglesias et al., 2023). After removing duplicates, we excluded primers with missing direction information, ambiguous bases, or adapter sequences, as well as those already present in the built-in primer dictionary for the same direction. Primers that yielded fewer than 500 distinct sequences were also excluded, leaving 272 primers for testing. For each primer, we sampled 500 distinct sequences and represented each sequence 20 times, giving 10,000 reads per library. Three replicate libraries were constructed per primer, producing 816 libraries in total.

Each library was tested with and without the built-in primer dictionary, using the same three window lengths and six thresholds as in Benchmark 1. This gave 36 conditions per library and 29,376 tests in total. The correct trimming endpoint was defined as the end of the primer. We calculated primer detection rates and the proportions of tests resulting in correct trimming, incomplete primer removal, or removal of biological bases. For each window length and dictionary condition, each primer’s detection rate was calculated across three replicates and six thresholds, giving 18 tests per primer.

3.2 Results

Benchmark 2 evaluated primer detection using 272 external primers, and three replicate libraries per primer, which yielded a total of 816 primers x libraries tests. Each library contained 10,000 reads from 500 unique sequences. Detection rates (Figure S3 A) were assessed using three primer length windows (15, 20, and 25 bp, Figure S3B), and six different fold-thresholds, both with and without the built-in primer database.

The tested primers ranged from 10 to 26 nucleotides in length, with a median of 19 nucleotides (Figure S3B). Most were 15–20 nucleotides long (236/272, 86.76%), and 20 nucleotides was the most common length (63/272, 23.16%).

Our results show that mean detection rates decreased as the length of the detection window increased (Figure S3 A). Across primers, replicates, and fold-thresholds, the 15bp window achieved a detection rate of 98.16%, both with and without the built-in primer database. At 20 bp, detection rates were 92.22% and 88.58% with and without the built-in primer database, respectively. When the primer sequence length was large (i.e., 25 bp window), the primer detection rate decreased dramatically, sometimes even below 60% (Figure S3A).

By using higher fold-threshold, we generally detected higher primer detection rates across all length windows. This improvement however depended on the sequence length and primer database condition. At 15 bp, detection increased from 92.28% (fold < 4) to 100% (fold < 64 or fold < 128), both with and without the built-in primer database. At 20 bp, the combination of a fold < 16 and the built-in primer database supports a 94.49% detection rate, while without the database the detection rate is of 92.89%. At 25 bp, increasing the threshold from 4 to 128 raised detection from 63.97% to 93.26% with the built-in primer database, and from 14.58% to 88.48% without it (Figure S3).

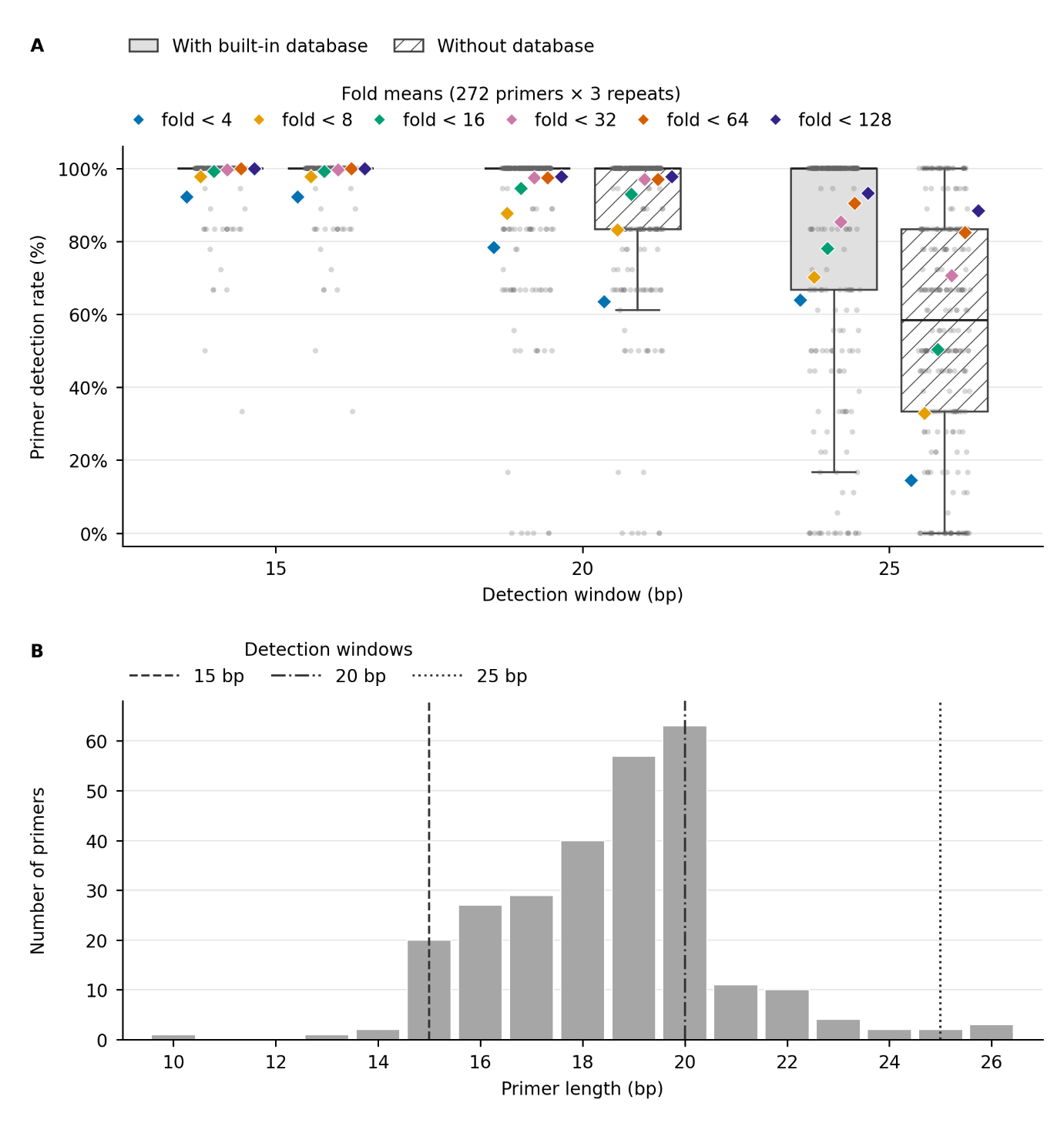

**Figure S3. Primer detection rate and primer length distribution during Benchmark 2.** (A) Primer detection rates using 15, 20, and 25 bp detection windows, either using the built-in primers database (gray boxes) or without the database (hatched boxes). Boxes show interquartile ranges, horizontal lines indicate medians. Gray points represent the individual detection rates of 272 primers, as the average of three replicates, and using six different fold thresholds (<4, <8, <16, <32, <64, and <128). Colored diamonds show the average primer detection rates at each of the six fold-thresholds, calculated across 272 primers and three replicates (816 libraries). (B) Length distribution of the 272 primers tested in Benchmark 2, with vertical lines delimitating the three detection length windows.

4. Benchmark 3: Measuring sequence loss during filtering

4.1 Methods

Public Illumina 16S V4 sequencing data were collected from 15 studies, including three studies for each of five different environments: human oral, aquatic, soil, animal gut, and human gut. Using Qiita sample information and ENA/SRA sequencing records, we excluded technical controls and duplicated data from any of the original studies, and selected the 20 samples with the highest reported read counts in each study, resulting in 300 different samples.

All samples were processed in a unique Meta2Data run using the VSEARCH approach. Samples were treated for quality filtering, primer trimming, removal of sequencing errors and chimeras, and grouping of the sequences with 97% identity into Operational taxonomic units (OTUs). The output OTU table provides a read count for each OTU for each sample included in the analysis, as well as a representative sequence for each OTU. Greengenes2 reference sequences were used to orient all OTU sequences consistently. A Greengenes2 classifier was also used to assign taxonomic labels to all OTU sequences. SEPP then placed the representative OTU sequences into the Greengenes 13_8 reference tree. The OTUs that could not be oriented or placed on the reference tree were removed from the analysis. Sequence read counts were recorded before orientation, after orientation, and after tree placement and filtering.

For each sample and study, we reported the proportions of OTUs and reads retained at each filtering step and across both steps combined. Retention at each filtering step was calculated as the amount remaining divided by the amount entering that step. Overall retention used the counts after merging but before orientation as the starting point. We checked that retained and removed reads summed to the input count. We also examined the sequence lengths and the best BLAST hits of each OTU sequence. Bray-Curtis dissimilarity was calculated to evaluate how these filtering steps affected the final community profiles. With these results, we showed where sequence losses occurred and how much of each sample was affected.

4.2 Results

Benchmark 3 assessed the effect of AmpliconTAXA filtering on the final output. AmpliconTAXA started with 48,103,128 reads from the merged PIP output. Orientation filtering removed 503,368 reads (1.046%), leaving 47,599,760 reads, while subsequent SEPP filtering caused no additional read loss in any sample (Figure S4A). The median sample-level orientation loss was 0.58%, although losses varied considerably among samples. The largest losses reached 83.82% in an animal-gut sample from PRJEB13147 and 40.37% in a soil sample from PRJEB15045. Bray-Curtis dissimilarities generally changed only marginally after filtering (Figure S4B). Among the 2,850 within-project pairs, the mean absolute change was 0.00416, and 91.09% of pairs changed by less than 0.01.

The orientation step excluded 2,436 features because their orientation could not be assigned to the correct direction. BLAST results for these excluded sequences against GG2 2024.09 revealed heterogeneous sequence similarity among the features (Figure S4C). We defined identity ≥90% and query coverage ≥80% as strong evidence of a match to the reference database. Among the excluded sequences, 54.51% of reads were assigned to features that met neither threshold or had no hit. In contrast, 2.64% of orientation-lost reads belonged to features meeting both the identity ≥90% and query coverage ≥80% thresholds. A further 28.48% met only the identity threshold, whereas 14.37% met only the coverage threshold.

The composition of removed reads differed among projects. In PRJEB15045, the identity-only category predominated, accounting for 61.78% of orientation-lost reads, while another 22.27% belonged to the coverage-only category. In contrast, 90.75% of removed reads in PRJEB13147 met neither threshold or had no reported match. Further examination of coverage, identity, and sequence length showed distinct characteristics among sequences meeting only one threshold (Figure S4D and F). The 148 identity-only features ranged from 61 to 373 nt in length and had a median query coverage of 30.5%, indicating that their high-identity matches generally covered only a limited portion of the full sequence. The 383 coverage-only features ranged from 61 to 275 nt and had a median identity of 80.15%, despite a median query coverage of 99%. Thus, orientation failures included both sequences with localized high-identity matches and sequences with extensive but lower-identity alignments to the reference. Despite these outliers, changes in beta diversity were generally minor, indicating that filtering had little effect on overall community composition. Samples with exceptionally high read loss likely represent low-quality or problematic datasets and should be carefully evaluated, and potentially excluded, in downstream analyses.

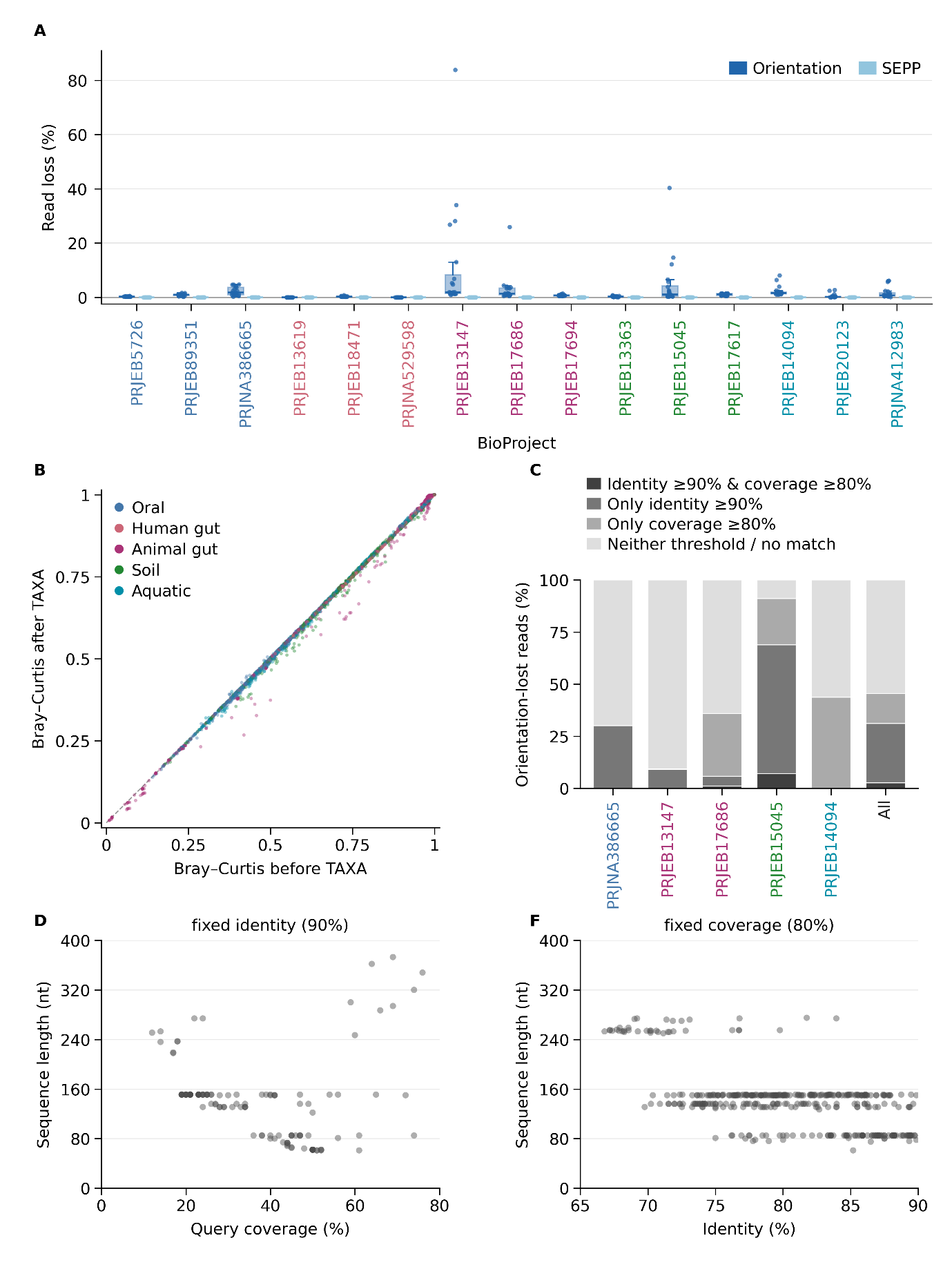

**Figure S4. Effects of AmpliconTAXA filtering in Benchmark 3.** (A) Read loss during orientation and SEPP filtering across 300 samples from 15 BioProjects covering five habitats (20 samples per project). Percentages are calculated relative to reads count before each step. Boxes show medians and interquartile ranges and dots represent individual samples. (B) Bray–Curtis dissimilarities before and after TAXA filtering for 8,850 within-habitat sample pairs, colored by habitat. The dashed line indicates equality. (C) Percentages of orientation-removed reads classified by BLAST against GG2 2024.09 (E ≤ 10⁻⁵), using identity ≥90% and query coverage ≥80%, both thresholds, identity only, coverage only, or neither/no match. “All” pools read counts across 15 BioProjects. (D) Sequence length versus query coverage for features meeting only the identity threshold. (F) Sequence length versus identity for features meeting only the coverage threshold. Each point in D and F represents one orientation-removed feature.

5. Benchmark 4: Measuring processing time and resource use

5.1 Methods

To assess how the computational cost of Meta2Data scales with sample number, 300 distinct BioSamples were selected from each of three projects: PRJNA529598, PRJEB18471, and PRJEB20123. Each sample was uniformly subsampled to 20,000 fragments, defined as read pairs for paired-end data and individual reads for single-end data. Nested sample subsets were used to maintain consistent input depth and sample sources across dataset sizes.

AmpliconPIP was run on subsets of 10, 25, 50, 100, 200, and 300 samples from PRJNA529598 using a fixed VSEARCH configuration and eight CPUs. AmpliconTAXA inputs were constructed by selecting equal numbers of samples from the 300-sample PIP outputs of the three projects, yielding total sample sizes of 30, 75, 150, 300, 600, and 900. At each size, SEPP tree-building and no-tree modes were compared using 20 CPUs.

Each condition was run independently once, recording elapsed time, actual CPU time, and whole-step peak memory. Measurements covered the complete module invocation but excluded data downloading, subsampling, and input preparation. TAXA measurements also excluded upstream PIP processing. Following output validation, computational scaling was assessed descriptively by plotting resource consumption against sample number separately for PIP and TAXA.

5.2 Results

AmpliconPIP runtime and CPU consumption increased steadily with sample number. Increasing the input from 10 to 300 samples raised runtime from 2.1 to 42.0 min and CPU consumption from 0.05 to 1.23 CPU-hours on the Roihu server at the CSC facilities, Helsinki, Finland. Peak memory increased overall from 1.49 to 6.77 GiB.

AmpliconTAXA showed different scaling patterns depending on whether tree construction was included. When using SEPP, increasing the input from 30 to 900 samples raised runtime from 18.8 to 42.9 min, and the CPU consumption from 2.28 to 8.53 CPU-hours. Without tree construction, runtime remained between 3.6 and 4.3 min, and CPU consumption ranged from 0.15 to 0.21 CPU-hours across all sample sizes. When using 900 samples, omitting tree construction reduced runtime by 90.2% and CPU consumption by 97.6%. However, peak memory remained comparable between the two modes, ranging between 41.7 and 50.8 GiB with SEPP, and between 46.2 and 49.8 GiB without tree construction.

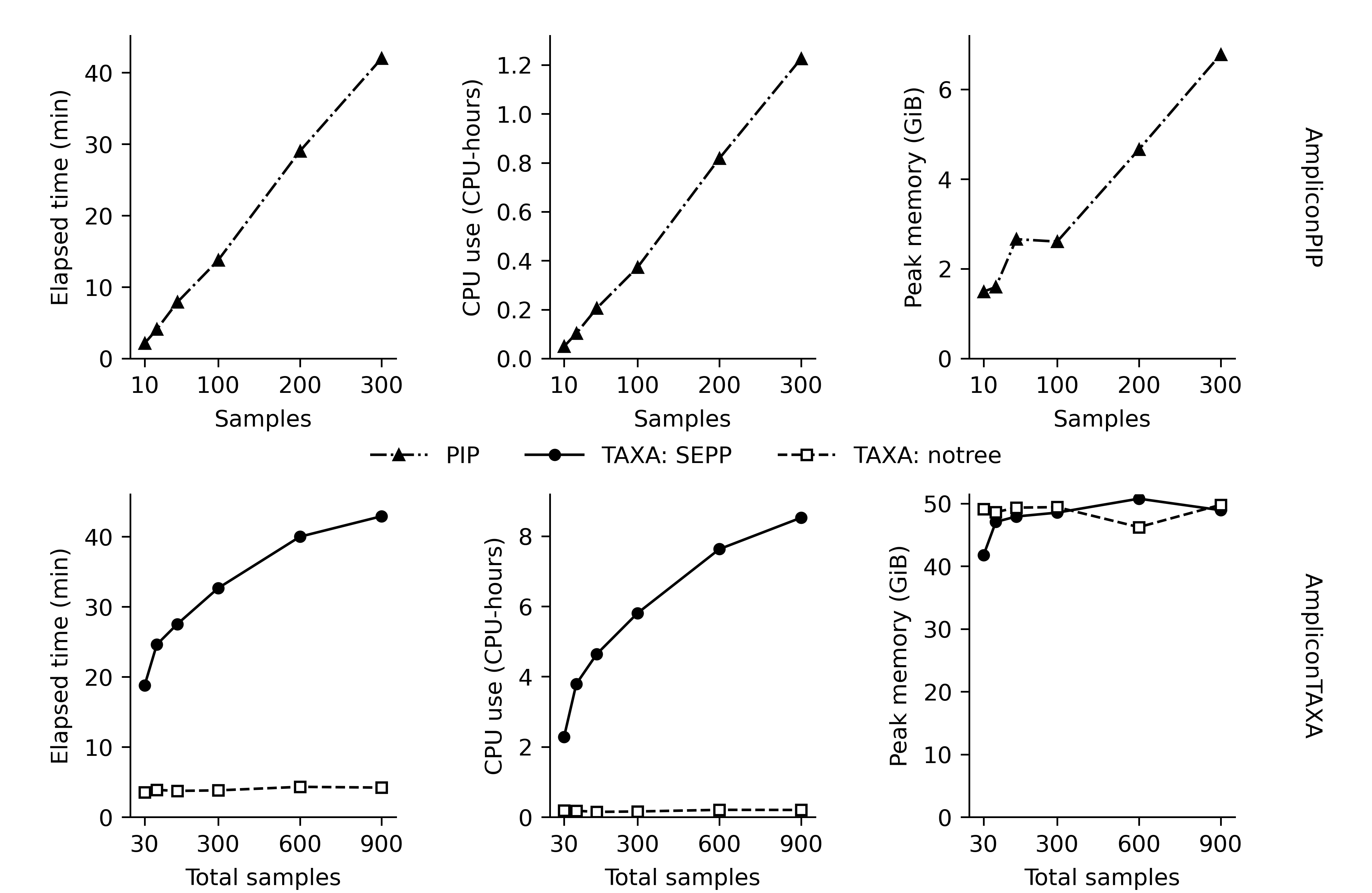

**Figure S5. Runtime and resource use during the AmpliconPIP and AmpliconTAXA modules.** The top row shows PIP; the bottom row compares TAXA with SEPP and without SEPP. Columns show elapsed time, CPU-hours, and peak memory, including file cache. Each condition was run once.
